# Modeling Laryngeal Dystonia through Spectral Analyses of Vocalizations in a Dystonia Mouse Model

**DOI:** 10.1101/2025.07.07.663183

**Authors:** Austin L. Fitzgerald, Jonathan A. Coello, Alyssa M. Lyon, Breanne L. Dao, Meike E. van der Heijden

## Abstract

Laryngeal dystonia is a task-specific, focal dystonia that disrupts vocal-motor control and significantly alters quality of life through impaired communication. Despite its early onset in many hereditary dystonias, effective treatments remain limited, in part due to the lack of a preclinical model that captures its circuit-level pathophysiology. Our experiment evaluates ultrasonic vocalizations (USVs) in *Ptf1a^Cre/+^;Vglut2^fl/fl^* mice, a cerebellum-specific generalized dystonia model, to assess translational relevance for laryngeal dystonia. At postnatal day 9, mutant mice demonstrated statistically significant reductions in total USV count, relative count of complex calls, and key spectral parameters— especially frequency modulation and power—mirroring phonatory abnormalities seen in human patients. Cluster analyses further revealed impaired vocal burst initiation, suggesting disrupted cerebellar coordination of temporal vocal-motor output. These findings support the model’s construct and face validity for cerebellar contributions to disordered phonation. By revealing these potential translational biomarkers, our study establishes a foundational platform for future mechanistic and interventional research in laryngeal dystonia.

**SUMMARY STATEMENT:** We demonstrate that a cerebellar general dystonia mouse model recapitulates vocal-motor deficits seen in laryngeal dystonia, providing a foundational platform for mechanistic studies and translational therapeutic discovery.

## INTRODUCTION

Laryngeal dystonia, also known as spasmodic dysphonia, is a rare neurological voice disorder that is a subcategory of focal dystonia (involuntary muscle contractions or abnormal movements affecting a specific body part or localized group of muscles). It is characterized by involuntary laryngeal muscle spasms that occur during speech, resulting in voice breaks, strained or strangled voice quality, or intermittent breathiness (Blitzer & Kohli, 2023). These symptoms significantly interfere with communication and reduce quality of life for affected individuals. Laryngeal dystonia has an estimated global prevalence of 1-14 per 100,000 with an average accurate diagnosis delays of 5.5 years (Hyodo et al., 2021; de Lima Xavier & Simonyan, 2019). Standard botulinum toxin injection treatment yields varied and impermanent results, with about a third of patients reporting no symptom improvement, underscoring the need for better disease-modifying approaches (Liu et al., 2024; Benninger & Smith, 2015).

Despite being classified as a focal dystonia, laryngeal dystonia shares circuit-level features with broader dystonic syndromes, including disrupted motor planning, abnormal sensorimotor integration, and impaired inhibition across varying cortical networks. Laryngeal dystonia often emerges early in broader genetic dystonic syndromes, especially in inherited or early-onset cases. In some DYT1 (THOR1A) families, cranial or cervical symptoms have been documented in childhood and can precede limb involvement, although this remains much less common than limb-first presentations (Bressman et al., 2000; Ozelius & Lubarr, 2016). In DYT6 (THAP1) dystonia, early laryngeal involvement is typical and often dominates the initial phenotype (Djarmati et al., 2009). Outcome-series further suggests that THAP1-positive patients achieve more variable benefit from global pallidus internus deep brain stimulation (GPi-DBS) than TOR1A carriers (Tisch & Kumar, 2021). These patterns support the idea that laryngeal dystonia arises from shared motor circuit dysfunction, manifesting in a focal, task-specific form. Studying cerebellar contributions to vocal control in this model may offer insight into early disease progression and highlight therapeutic windows missed by limb-focused models.

Treatment options for laryngeal dystonia are limited. Botulinum toxin injections are the standard intervention, but effects are temporary, responses vary, and side effects like breathiness or difficulty swallowing are common. Voice therapy and oral medications offer inconsistent relief. Clinical series also report varied and sometimes absent improvement of speech after pallidal deep brain stimulation (DBS), highlighting the need for circuit-targeted strategies specific to laryngeal dystonia (Artusi et al., 2020; Kupsch et al., 2006). These limitations reflect a gap in our understanding of the disorder’s neurobiological underpinnings, especially at the circuit level, that must be addressed to develop better treatments.

Neuroimaging has identified both structural and functional alterations in multiple areas involved in motor control, such as the basal ganglia, thalamus, sensorimotor cortex, supplementary motor area, and cerebellum (Kshatriya et al., 2024). A recent meta-analysis compiling data from over 500 patients with laryngeal dystonia showed consistent abnormalities in motor and sensorimotor regions of the brain, supporting the view that the disorder involves not just the larynx itself but the neural circuits responsible for planning and coordinating voice production (Kshatriya et al., 2024). One region that has received increasing attention in dystonia research is the cerebellum. Traditionally associated with coordination, balance, and motor learning, the cerebellum is now better understood to play important roles in speech timing and fine motor control of the vocal apparatus. Functional imaging studies demonstrate hyper-activation of the cerebellum during symptomatic speech in laryngeal dystonia, supporting a cerebellar contribution to disordered vocal motor control (Simonyan & Ludlow, 2010). These cerebellar disruptions may help explain why patients with laryngeal dystonia struggle to regulate pitch and vocal intensity during speech. In fact, human laryngeal dystonia features aberrant acoustic events including exaggerated pitch-shift reflexes and impaired auditory-motor adaptation, reflecting a hyper-reactive yet poorly calibrated feedback loop (Dwenger et al., 2025; Edgar et al., 2001; Kim & Larson, 2019; Larson & Robin, 2016; Ludlow et al., 2018; Sapienza et al., 1999).

A major obstacle to developing better treatments for laryngeal dystonia is the lack of an animal model that captures its task-specific vocal-motor features (Roy et al., 2024). Most existing rodent models of dystonia are primarily striatal or cortical and recapitulate generalized or limb motor dysfunction but lack any voice-specific phenotype, limiting their utility for studying laryngeal symptoms or circuitry (Arriaga & Jarvis, 2013; Hipolito et al., 2023). This gap has slowed efforts to identify circuit-specific biomarkers and test interventions translatable to vocal control. However, mouse ultrasonic vocalizations (USVs) offer a promising solution. While mice do not produce speech like people, their USVs are high-frequency, structured vocal signals that require precise coordination of laryngeal and respiratory muscles. These calls rely on intact cerebellar and auditory feedback loops, making them a surprisingly rich analog for vocal-motor function (Lahvis et al., 2011). USVs can be quantitatively analyzed for call type, frequency, modulation, power, and duration—features that may parallel the disruptions seen in patients with laryngeal dystonia (Marchese et al., 2024).

Our study uses a well-characterized mouse model of generalized dystonia driven by cerebellum-specific circuit disruption, the *Ptf1a^Cre/+^;Vglut2^fl/fl^*, mouse model. This model replicates dystonic movements that improve with cerebellar-targeted DBS and meets key standards for animal model validity: (1) face validity—visible muscle contractions similar to those in patients; (2) construct validity—abnormal cerebellar function consistent with human and rodent dystonia; and (3) predictive validity—postural and limb symptoms respond to cerebellar DBS, paralleling patient outcomes (Brown et al., 2022; White & Sillitoe, 2017). By analyzing USVs in dystonic versus control mice, this study aims to validate this circuit-level model for laryngeal dystonia. This study focused on postnatal day 9 (P9) mice due to this time point being a developmental stage where cerebellar circuits are actively maturing and USVs are reliably produced (Altman & Bayer, 1997; Yin et al., 2016).

In this manuscript, we set out to investigate whether mice with cerebellar dysfunction-related generalized dystonia will show mechanical USV abnormalities mimicking human laryngeal dystonia. This work addresses a critical unmet need in the field: the development of an animal model that links neural circuit dysfunction to vocal-motor abnormalities with preclinical relevance to human disease.

## RESULTS

### USV Call Types Show Quantitative Reductions in Cerebellar-Dysfunction General Dystonia Mutant Mice at age P9

First, to determine whether *Ptf1a^Cre/+^;Vglut2^fl/fl^* mutant mice display altered USV patterns compared to *Vglut2^fl/fl^* controls, we categorized calls emitted by P9 pups into call types based on shape, modulation, and syllable number using Metris Sonotrack software **(Fig. 1).** We found that all call types were emitted by both mutant and control pups.

**Figure 1.**
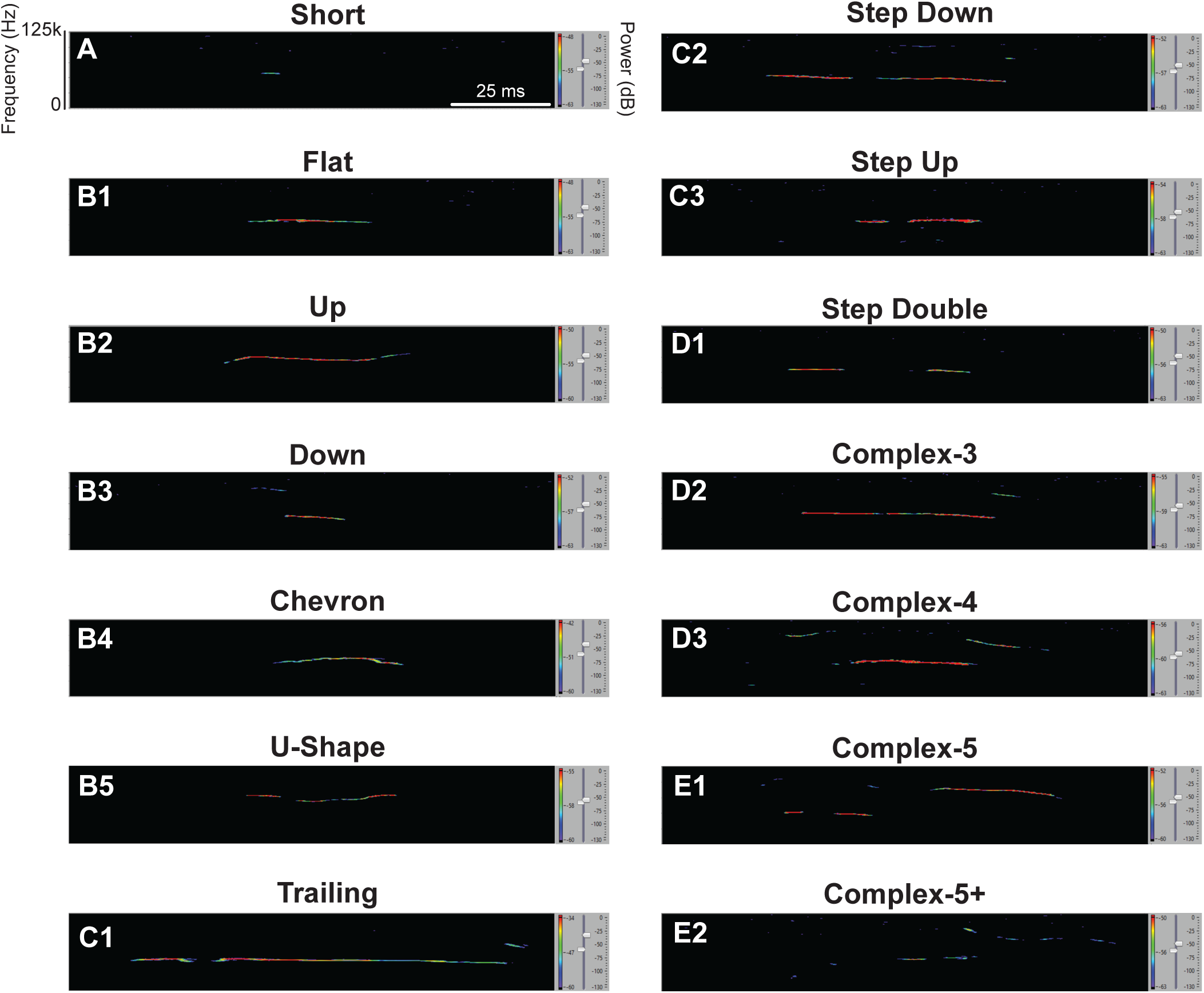
Example spectrograms of categorized USV call types recorded from P9 mouse pups. Each panel shows an example USV spectrogram annotated by the Metris Sonotrack software according to spectral features which include frequency (Y-axis, kHz), time (X-axis, ms), and power intensity (color scale, dB). Calls are categorized based on frequency modulation, shape, and complexity. These spectrograms are intended as qualitative examples only. All USV call types were categorized according to Metris Sonotrack Call Classification which is based upon call classification principles described in Portfors (2007). Additionally, ‘syllables’ refer to individual vocal elements within a call and are synonymous with Sonotrack’s definition of ‘elements’ (Portfors 2007): (A) Short Calls are brief, simple calls with short duration <15 ms. These calls are often isolated with little to no modulation. One Syllable Calls include: (B1) Flat Calls are calls with a nearly constant frequency throughout. This indicates no major pitch change, producing a monotone tone. (B2) Up Calls where the frequency increases over time corresponding to a rising pitch. (B3) Down Calls where the frequency decreases over time corresponding to a falling pitch. (B4) Chevron Calls are V- or inverted V-shaped calls with a sharp rise and fall (or fall and rise) in frequency. They are often brief and energetic. (B5) U-Shape Calls are a descending then ascending smooth curve in frequency. Two Syllable Calls include: (C1) Trailing Calls end with a long, fading frequency tail. (C2) Step Down Calls have one or more abrupt drops in frequency and resemble a stair-step downward. (C3) Step Up Calls abruptly shift upwards in frequency like stair-steps going upward. 3-4 Syllable Calls include: (D1) Step Double Calls are composed of two sequential step-like changes in frequency, creating a double-stepped shape. (D2) Complex-3 Calls contain three distinct modulations or frequency shifts and suggest more sophisticated communication. (D3) Complex-4 Calls are four-part calls with varying pitch directions or shapes, reflecting more nuanced or emotionally complex vocalizations. 5+ Syllable Calls include: (E1) Complex-5 Calls are five-segment calls with multiple frequency modulations. They are typically rare. (E2) Complex-5+ Calls are calls with more than five modulated components which make them highly elaborate.

Total number and proportional distribution of calls across syllable-defined categories were compared between groups **(Fig. 2).** Mutant mice emitted statistically significantly fewer total USVs compared to controls (*p*=0.0005) (**Fig. 2A**). Additionally, all call type categories—Short, One-, Two-, Three to Four-, and Five+ syllable calls—were statistically significantly reduced in absolute number of calls in mutant mice pups (*p*=0.0001; *p*=0.0020; *p*=0.0055; *p*=0.0068; *p*=0.0060, respectively) **(Fig. 2B1,C1,D1,E1,F1).** These findings suggest that cerebellar dysfunction in this dystonia model not only reduces overall vocal output but also disrupts the production of spectrally and temporally complex vocalizations, indicating impaired initiation or coordination of vocal-motor patterns early in development.

**Figure 2:**
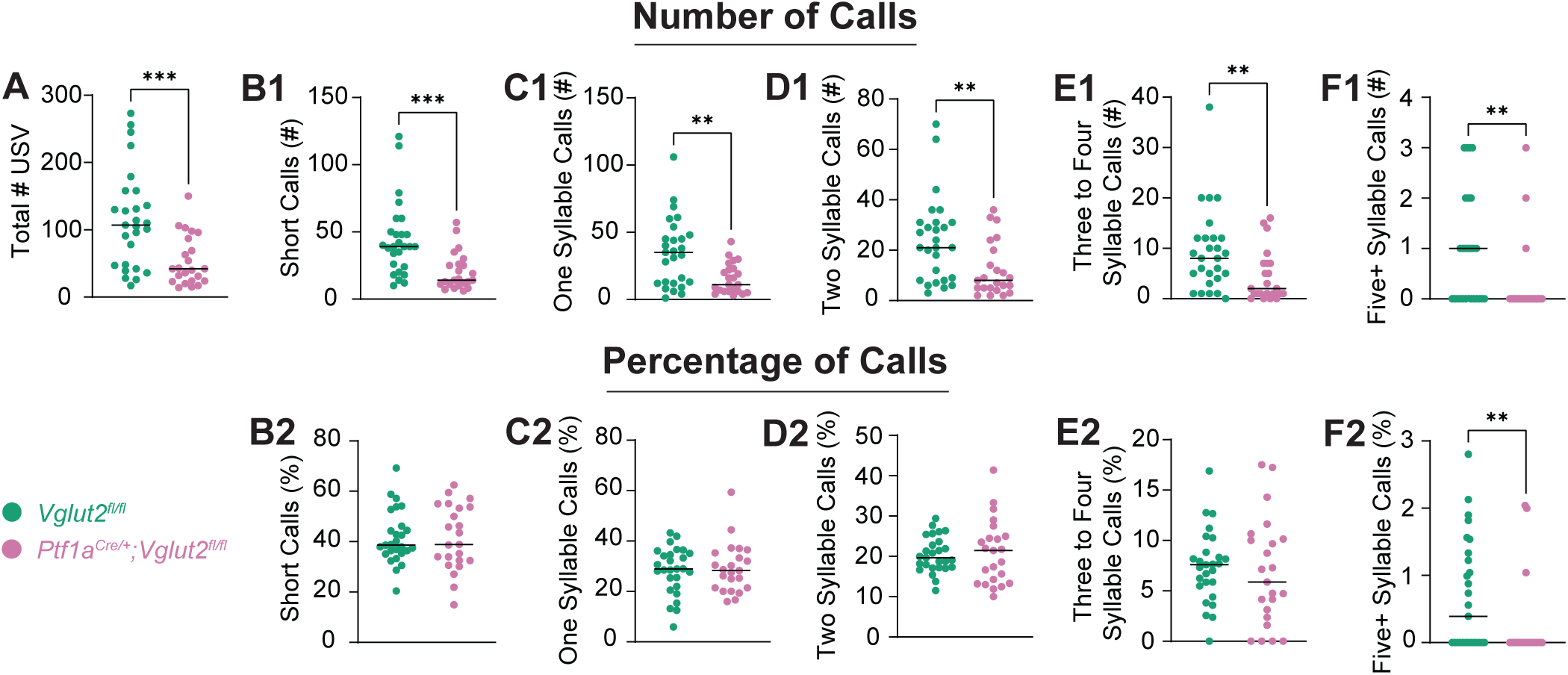
Quantitative analysis of USV call types reveals fewer total vocalizations and a statistically significant reduced percentage of the most complex calls in dystonic pups. Graphs represent individual data points and show group means for total USV count and call type distributions grouped by syllable number. Individual data points represent biological replicates (one pup; *n*=27 *Vglut2^fl/fl^* controls, 23 *Ptf1a^Cre/+^;Vglut2^fl/fl^* mutants; experiment performed once per pup). (A) Total number of USVs emitted per pup. (B1, C1, D1, E1, F1) Number of Short, One-, Two-, Three to Four-, and Five+ syllable calls emitted per pup, respectively. (B2, C2, D2, E2, F2) Percentage of Short, One-, Two-, Three to Four-, and Five+ syllable calls emitted per pup, respectively. Syllable categorization of call types is explained in Figure 1. Statistical comparisons were performed using unpaired, two-tailed Mann-Whitney U tests. Statistically significant reductions were observed in total number of USVs, absolute numbers of all call type categories, and relative Five+ syllable calls in mutants compared to controls. Data suggests a marked reduction in both number and complexity of vocalizations in mutant animals. \**p*<0.05; \*\**p*<0.01; *\*\*\*p*<0.001.

Next, we set out to investigate whether the relative proportion of call types was different between mutant and control pups. Only the percentage of the most complex vocalizations, Five+ Syllable Calls, was statistically significantly reduced in mutants (*p*=0.0089) suggesting disproportionate reductions in complex call production, even if Three to Four Syllable Calls do not show a statistically significant reduction. **(Fig. 2E2,F2).** All other percentage categories did not differ in a statistically significant manner between groups. This aligns with our hypothesis that while number of calls will uniformly be reduced in mutant mice compared to controls, the percentage of complex calls only will be reduced in mutants. Ultimately, this result suggests that while overall vocal output is reduced, mutants demonstrate an additional relative loss of more complex call types. This is indicative of a disrupted and/or less precise audio feedback loop given the fact that complex calls require precise audio feedback modulation. These findings confirm that core spectral features broadly are disrupted at the critical development stage and preliminarily establish this general dystonia mouse model’s potential for accurately capturing cerebellar vocal-motor deficits.

### Short Call Acoustic Parameters Are Altered in Mutants

We next examined pertinent acoustic features of Short Calls (<15 ms duration), which was consistently the predominant call category. Multiple parameters including duration, frequency, power, and frequency modulation were analyzed per pup and compared between groups **(Fig. 3).** Based on human literature, we expected group differences between control and dystonic mice to be most pronounced in frequency modulation and power as these features correspond to impaired laryngeal muscle control and audio-vocal feedback seen in laryngeal dystonia with significantly less to no statistically significant differences in average vocal frequency parameters as a simple change in tonality of voice alone is not a distinct feature of laryngeal dystonia (Kshatriya et al., 2024; Marchese et al., 2024).

**Figure 3:**
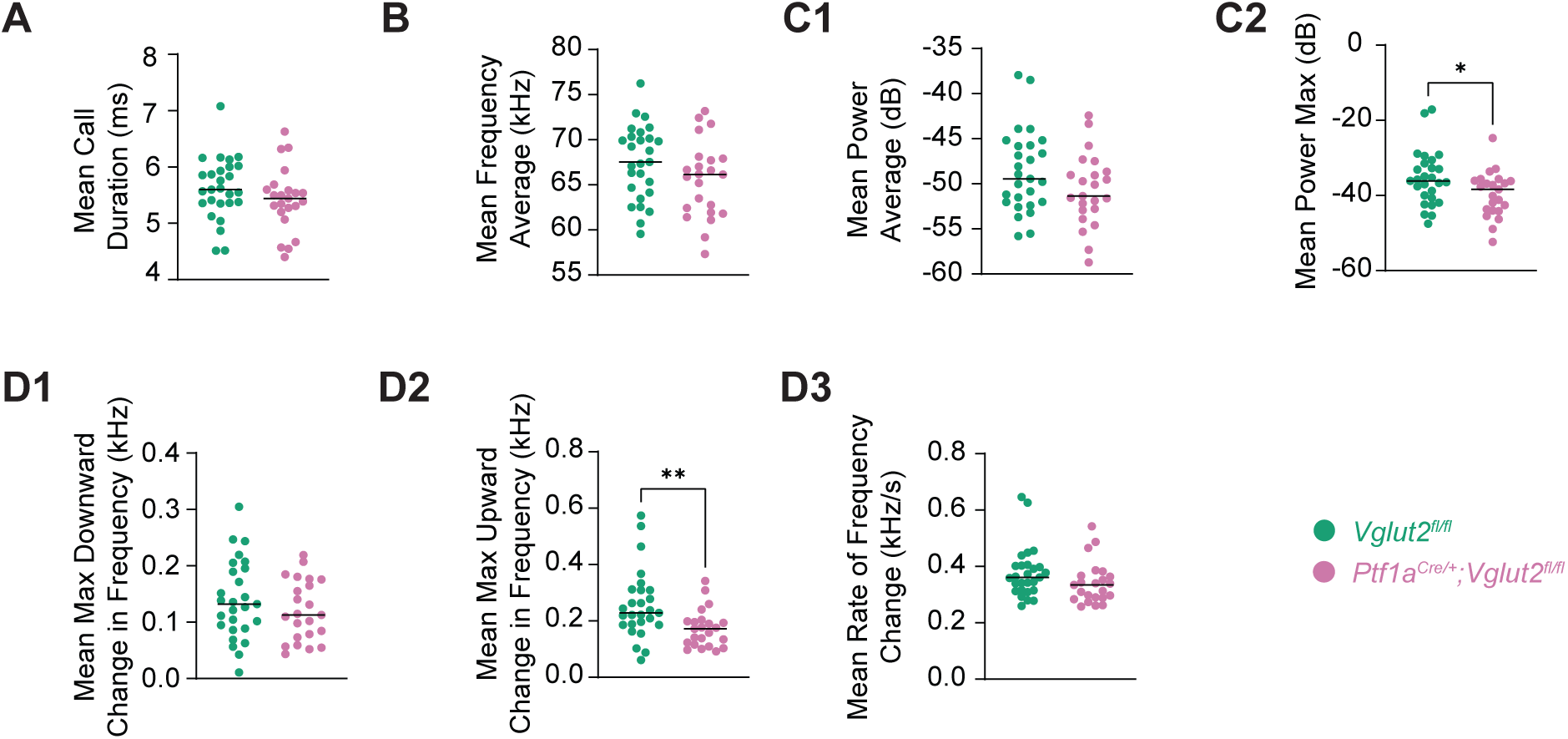
Analysis of acoustic properties of short calls indicates altered signal structure in dystonic pups. Graphs represent individual data points and show group means for various acoustic parameters of short calls (<15 ms in duration). Individual data points represent biological replicates (one pup; *n*=27 *Vglut2^fl/fl^* controls, 23 *Ptf1a^Cre/+^;Vglut2^fl/fl^* mutants; experiment performed once per pup). (A) Mean short call duration (ms). (B) Mean frequency average (kHz). (C1, C2) Relevant power parameters of mean power average (dB) and mean power maximum (dB), respectively. (D1-D3) Frequency modulation parameters of mean maximum downward change in frequency (kHz), mean maximum upward change in frequency (kHz), and mean rate of frequency change (kHz s^-1^), respectively. Statistical comparisons were performed using unpaired, two-tailed Mann-Whitney U tests. Statistically significant differences were detected in upward frequency change and maximum power, indicating acoustic alterations in short calls emitted by mutant pups. \**p*<0.05; \*\**p*<0.01; *\*\*\*p*<0.001.

Compared to controls, mutant pups showed a statistically significant reduction in maximum power (*p*=0.0347) and maximum upward change in frequency modulation (*p*=0.0031) **(Fig. 3C2,D2).** Other parameters reported in **Figure 3** do not show statistically significant differences between the two groups. These findings indicate that mutants exhibit alterations in acoustic structure and strengthen the translational validity of our model since the vocal impairment parameters that show a statistically significant reduction align with the vocal impairments observed as a product of reduced laryngeal muscle coordination observed in laryngeal dystonia patients.

### Long Call Acoustic Parameters Are Also Altered in Mutants but More Limited

We separately analyzed Long Calls (>15 ms duration) for comparable acoustic parameters **(Figure 4)**. Among all variables tested, only mean rate of frequency change differed between groups in a statistically significant manner (*p*=0.0314) (**Fig. 4D3**). Other frequency modulation, duration, frequency, and power parameters did not show statistically significant differences. While this data suggests nominal acoustic changes in longer calls, potentially indicating that more complex vocalizations may be preserved in spectral quality despite being less frequent in mutants, the fact that a frequency modulation parameter did show a statistically significant reduction in mutant mice and it is a different one than what was present in short calls leads us to reasonably come to similar conclusions as our short call results—this result is a product of reduced laryngeal muscle coordination similar to those observed in human laryngeal dystonia patients.

**Figure 4:**
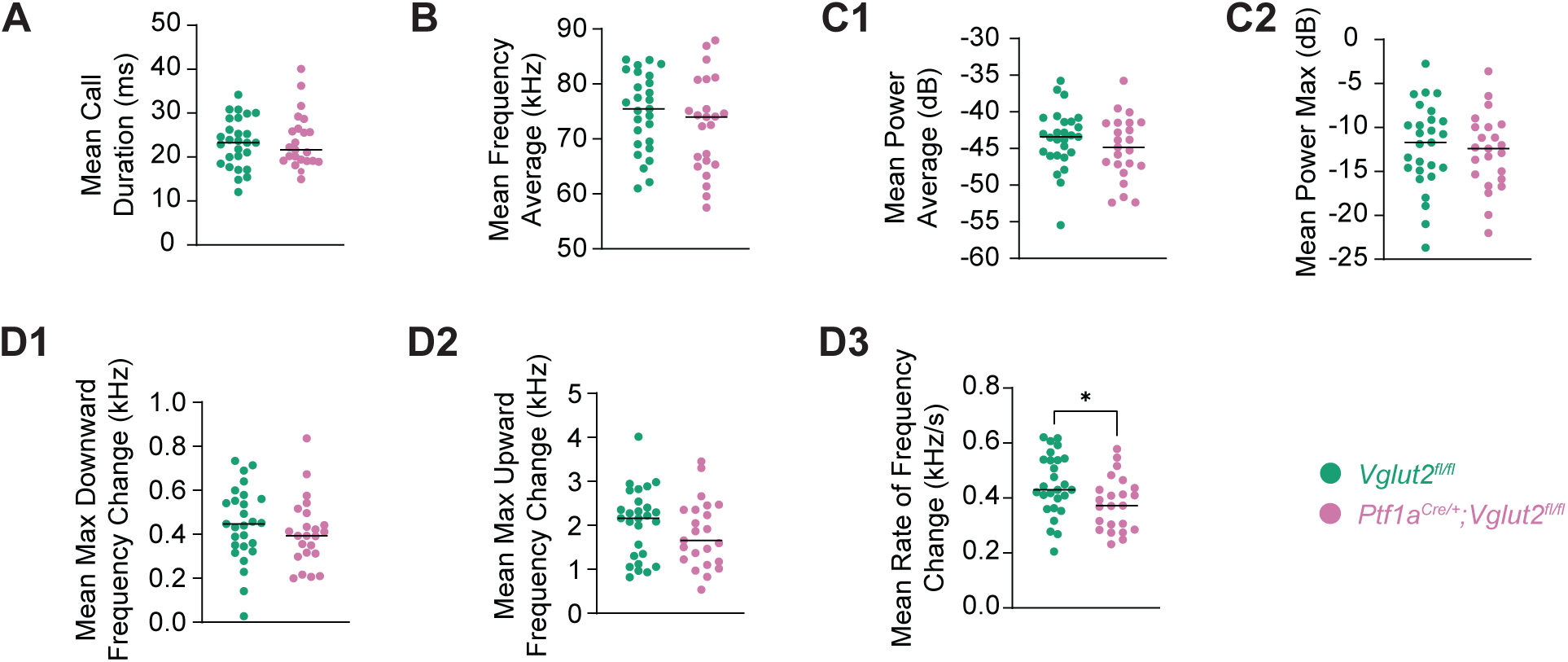
Analysis of acoustic properties of long calls shows altered signal structure in dystonic pups. Graphs represent individual data points and show group means for various acoustic parameters of long calls (>15 ms in duration). Individual data points represent biological replicates (one pup; *n*=27 *Vglut2^fl/fl^* controls, 23 *Ptf1a^Cre/+^;Vglut2^fl/fl^* mutants; experiment performed once per pup). (A) Mean long call duration (ms). (B) Mean frequency average (kHz). (C1, C2) Relevant power parameters of mean power average (dB) and mean power maximum (dB), respectively. (D1-D3) Frequency modulation parameters of mean maximum downward change in frequency (kHz), mean maximum upward change in frequency (kHz), and mean rate of frequency change (kHz s^-1^), respectively. Statistical comparisons were performed using unpaired, two-tailed Mann-Whitney U tests. A statistically significant difference was detected in mean rate of frequency change, indicating acoustic alterations in long calls emitted by mutant pups. \**p*<0.05; \*\**p*<0.01; *\*\*\*p*<0.001.

### Call Clustering Patterns Differ in Mutant Mice

Finally, to investigate if and how USV patterns evolve temporally between our two groups, we analyzed call clustering behavior across pups. We defined clusters as sequences of two or more unique calls in a row in which the start time of each call was within 0.5 seconds of the end time of the previous call. Inter-call interval is defined as the time between the end of a prior call and the start of the next call. This metric reflects the organization of vocal bursts which can potentially offer insight into vocal planning, effort, and audio-vocal feedback function. While average duration of clusters, average number of calls in clusters, and average cluster inter-call interval did not differ in a statistically significant way between groups **(Figs 5B-5D)**, mutant pups did exhibit a statistically significant reduction in the total number of clusters compared to controls (*p*=0.0001) **(Fig. 5A).** This finding suggests that while the structure of each vocal cluster may be preserved, mutant mice produce fewer discrete bouts of vocal activity overall. This finding aligns with our finding that mutant mice produce a statistically significant reduction in total number of USVs and seemingly simplified vocal behavior in these pups. This may reflect impairments in the initiation or coordination of vocal bursts in mutant mice.

**Figure 5:**
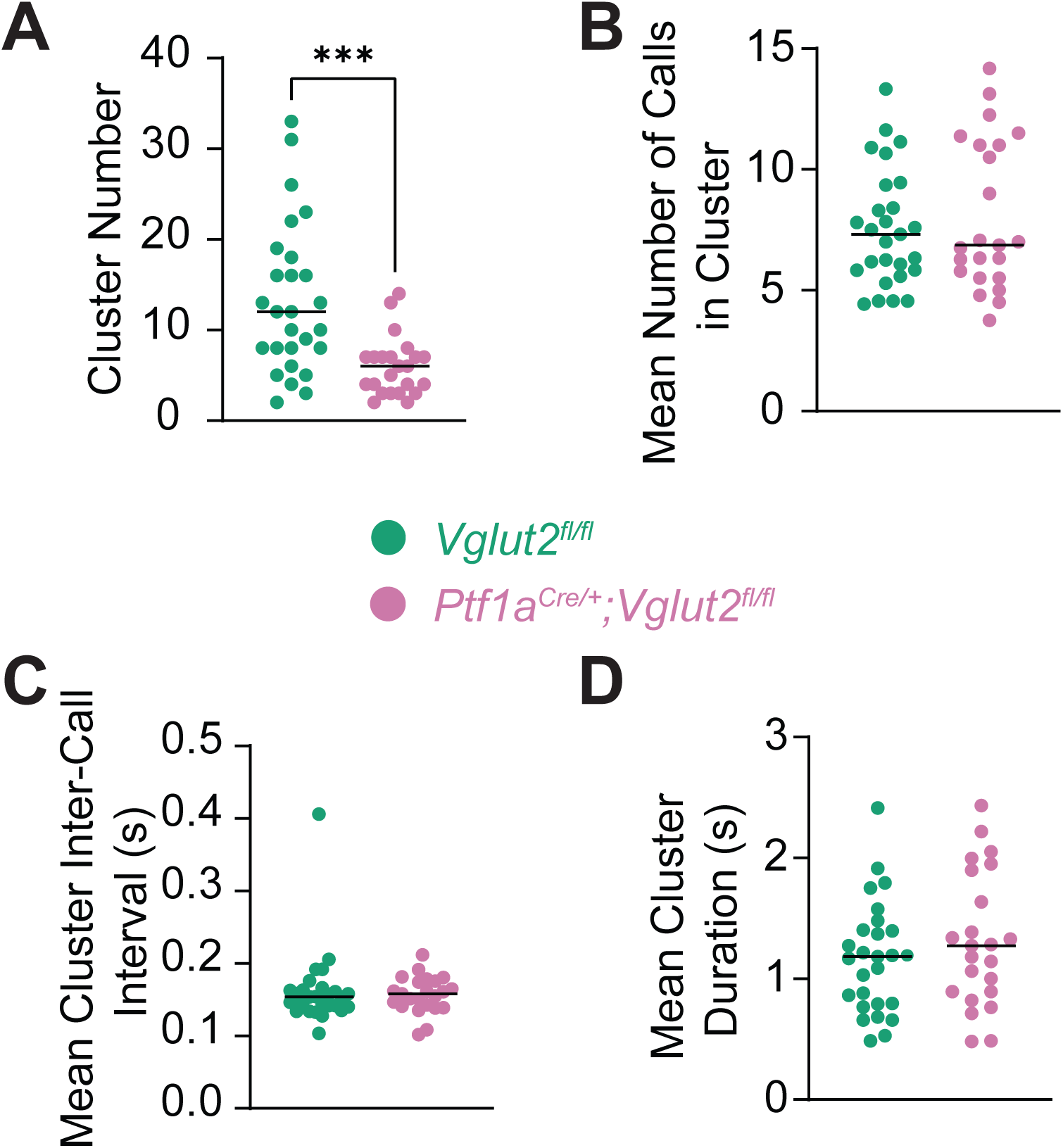
Cluster analysis of USV patterns reveals altered vocal grouping behavior in dystonic pups. Graphs represent individual data points and show group means for cluster-based metrics derived from USVs. Individual data points represent biological replicates (one pup; *n*=27 *Vglut2^fl/fl^* controls, 23 *Ptf1a^Cre/+^;Vglut2^fl/fl^* mutants; experiment performed once per pup). A cluster was defined as a group of 2 or more unique calls where the start time of each call occurred within 0.5 s of the end time of the previous call (this also defines our inter-call interval). (A) Number of detected USV clusters per pup. (B) Mean number of calls per cluster. (C) Mean cluster inter-call interval (s) within clusters. (D) Mean cluster duration (s). Statistical comparisons were performed using unpaired, two-tailed Mann-Whitney U tests. A statistically significant reduction in the number of USV clusters in mutant pups compared to controls was observed, suggesting altered temporal organization of vocalizations. \**p*<0.05; \*\**p*<0.01; *\*\*\*p*<0.001.

Overall, successfully identifying these specific differences in vocalization patterns is a promising step towards validating our *Ptf1a^Cre/+^;Vglut2^fl/fl^* mouse model as an adaptable model for laryngeal dystonia. In addition to the spectral and structural call abnormalities observed, our findings that mutant mice emitted a statistically significant fewer amount of temporal call clusters compared to controls suggest that dystonic pups produce fewer calls, simpler calls, and demonstrate altered organization of vocal output over time. These findings all support the idea that cerebellar dysfunction impacts not only vocal content but also how vocalizations are sequenced and structured across time, substantiating the use of this generalized dystonia model to study cerebellar contributions to vocal-motor coordination and audio-vocal feedback which are hallmark features impaired in human laryngeal dystonia. This lays foundational groundwork for further validation studies and future research into disease mechanisms and therapeutic strategies.

## DISCUSSION

### Cerebellar Dysfunction Disrupts Both the Complexity and Timing of Vocal-Motor Output in Neonatal Mice

Our findings highlight that cerebellar dysfunction alters not only the complexity but also the temporal architecture of USVs in our *Ptf1a^Cre/+^;Vglut2^fl/fl^* dystonic mouse model. These findings also align with prior work showing that the cerebellum acts as an integrative hub for temporal and spatial coordination of speech motor output (Simonyan & Fuertinger, 2015). Additionally, the observed statistically significant reduction in clustered vocal sequences suggest disrupted sequencing or initiation of vocal-motor bursts, supporting recent findings that implicate the cerebellum in temporal coordination and volitional control of speech in laryngeal dystonia (Kothare et al., 2022; Kshatriya et al., 2024).

Though mice do not produce human speech, their clustered USV emissions offer a behavioral analogue of structured phonation. Therefore, the pattern of our findings in this initial study are incipiently consistent with human studies of laryngeal dystonia in which patients exhibit impaired audio-vocal feedback control and delayed or fragmented speech initiation and are promising results towards establishing this mouse model as anovel animal model for laryngeal dystonia (Behroozmand & Sangtian, 2018; Rogić Vidaković et al., 2023; Weerathunge et al., 2022). Our cluster analysis revealed that while the structure within clusters (call number per cluster, average cluster duration, and average inter-call latency) were relatively preserved, the frequency of initiating discrete clusters was reduced in dystonic mice in a statistically significant manner. This strengthens the suggestion that cerebellar output is particularly essential for initiating and sustaining patterned vocal behaviors rather than modulating fine timing of each vocalization within a cluster.

In summary, these findings underscore the utility of this specific *Ptf1a^Cre/+^;Vglut2^fl/fl^* cerebellum-targeted generalized dystonia model in studying both spectral and temporal features of vocal-motor coordination which are also key components disrupted in human laryngeal dystonia. Importantly, this reinforces the need for future behavioral paradigms that test cerebellar contributions to vocalizations under stress or task-specific conditions, better approximating the demands of human speech (Marchese et al., 2024).

### Establishing Face Validity through Frequency Modulation and Call Complexity as Translational Biomarkers for Human Laryngeal Dystonia

The frequency modulation changes and reduction in complex call types we identified in mutant mice resemble observations in human laryngeal dystonia. Patients with laryngeal dystonia often demonstrate abnormal pitch control, exaggerated pitch-shift responses, and reduced adaptive responses to altered auditory feedback (Behroozmand & Sangtian, 2018; Thomas et al., 2021). Recapitulating features seen in human laryngeal dystonia patients, our dystonic mice showed statistically significant reductions in maximum upward frequency modulation in short calls and rate of frequency change in long calls. These parallels contribute to the relevance of our endpoints as potential behavioral biomarkers across species.

Furthermore, the observed loss of spectrally rich, complex calls in mutants resembles voice breaks documented behaviorally in human laryngeal dystonia, although direct cerebellar correlates in patient acoustic studies remain to be determined. Thus, the mouse USV system provides a simplified yet powerful platform for investigating circuit-level contributors to disordered phonation.

### Limitations and Future Directions

While our findings are promising, several limitations must be acknowledged. First, USVs are an indirect proxy for phonation and, while useful, do not capture the full scope of laryngeal biomechanics. Second, the generalized dystonia model may not fully replicate the task-specific nature of human laryngeal dystonia (Roy et al., 2024). Future experiments will potentially involve collaborations with voice centers and speech pathologists to bridge the behavioral phenotype with human vocal pathology more directly. For example, leveraging data from human laryngeal scope studies and pitch-control paradigms could strengthen translational parallels.

Our study focuses on postnatal day 9 developmentally. This is a timepoint at which pup vocalizations serve key communicative and regulatory roles. Given that human laryngeal dystonia can emerge in childhood but also adulthood, it is essential to determine whether the observed abnormalities persist in later developmental stages. A follow-up study of adult vocal behaviors will help to determine whether these early deficits persist into adulthood and provide insight into their progression and stability.

### A Platform for Therapeutic Screening

Ultimately, our cerebellum-specific conditional knockout mouse model opens the door to new opportunities for mechanistic and interventional studies for a predominant focal dystonia subtype. Additionally, laryngeal dystonia frequently emerges early in genetic forms of dystonia, such as DYT6, THAP1-associated dystonia, often preceding other motor symptoms. This early laryngeal involvement underscores its importance as a critical treatment target in genetically susceptible individuals (Artusi et al., 2020; Bressman et al., 2000; Djarmati et al., 2009; Kupsch et al., 2006; Ozelius & Lubarr, 2016). Transcranial magnetic stimulation (TMS), DBS, and pharmacological approaches could be applied in this model to test circuit reversibility or compensation—strategies that are actively being explored in human trials (Rogić Vidaković et al., 2023). By continuing to develop this mouse model in conjunction with human data, we move closer to a viable preclinical model for mechanistic exploration and to identify therapeutic targets in laryngeal dystonia.

## MATERIALS & METHODS

### Animal Models and Ethical Compliance

All procedures were approved by the Virginia Tech Institutional Animal Care and Use Committee (IACUC Protocol #23-181) and adhered to NIH guidelines for the care and use of laboratory animals. Experiments were performed on C57BL/6J mice, with the dystonic group comprised of *Ptf1a^Cre/+^;Vglut2^fl/fl^* mice (n=23) with glutamatergic neurotransmission selectively eliminated from the inferior olive to Purkinje cells (van der Heijden et al., 2022). Littermate *Vglut2^fl/fl^* mice served as controls (n=27). Both sexes were used, and all recordings were conducted at P9, a developmental timepoint when pups vocalize reliably following maternal separation and cerebellar circuits are undergoing active refinement (Altman & Bayer, 1997; Yin et al., 2016).

### Experimental Design and Recording Setup

Mouse pups (n=50 from 9 litters) were separated from the dam and immediately and individually placed in a sound-isolated Metris SmartChamber for 120 seconds of free vocalization. USVs were recorded using a Metris Gold Foil Electrostatic Transducer and digitized at a sampling rate of 250 kHz. Spectrograms were analyzed using Metris Sonotrack software (v1.4.7, Metris B.V., Netherlands).

This USV experiment was designed to assess cerebellar involvement in early vocal production by comparing vocalization quantity, acoustic structure, and temporal dynamics between dystonic and control pups. Nine animals emitting 10 or fewer calls during the 120 second recording period were excluded according to a pre-established criterion to ensure statistical reliability.

All mice from available, qualifying litters were recorded, and vocalizations were recorded in a consistent environment. Investigators were blinded to genotype during data collection.

### USV Parameter Definitions and Classification

Each USV call was categorized using Metris Sonotrack software based on spectral shape, duration, and modulation characteristics. Call types include short, flat, up, down, chevron, U-shape, trailing, step down, step up, step double, complex-3, complex-4, complex-5, and complex-5+ calls, among others, as defined in Portfors (2007). Individual local elements—termed “syllables” in this study—correspond to what Metris Sonotrack refers to as “elements” (Lahvis et al., 2011; Portfors, 2007). Sample spectrograms of call types can be found in **Figure 1**.

Acoustic parameters analyzed for each call from each pup included: (1) time-based features of call duration, start time, and end time; (2) frequency features of frequency start, end, minimum, maximum, and average; (3) power features of power average, power maximum, and power at frequency minimum, maximum, and average; and (4) frequency modulation features of rate of frequency change, maximum upward frequency change, and maximum downward frequency change.

Calls were additionally divided into short (<15 ms) and long (>15 ms) categories for each mouse, with separate acoustic means computed within these duration groups to minimize skew from call length distribution. In multi-syllabic calls, each syllable was independently analyzed.

### Cluster Analysis

USVs were grouped into temporal clusters, defined as at least 2 consecutive calls where the start of each call occurred within 0.5 seconds of the previous call’s end. Parameters extracted per cluster included: number of clusters per mouse pup, number of calls per cluster, mean inter-call interval, and cluster duration.

### Statistical Analysis

Averages of each acoustic parameter for each mouse were calculated in Microsoft Excel and cluster analysis parameters were extracted from the original Metris Sonotrack data set for each mouse utilizing MATLAB (Mathworks, United States). The remainder of the statistical analysis was conducted using GraphPad Prism (v10.5.0, GraphPad Software, San Diego, CA). Five more mice were excluded from the dataset after an outlier analysis. Group comparisons were performed using unpaired, two-tailed Mann-Whitney U tests due to the expected non-normal distribution of USV parameters. Spread of data was visualized via scatterplots to enhance interpretation **(Figs 2-5).**

A post-hoc power analysis was performed using G*Power (v3.1.9.6) for the primary endpoints of total USVs and maximum upward frequency modulation for short calls using Cohen’s d=1.0701 and 0.8522, respectively, and samples sizes of 27 (control) and 23 (mutant) estimated 95% and 82%, respectively, at α=0.05 (two-tailed), indicating adequate sensitivity. No power analysis was conducted for secondary comparisons due to the exploratory nature of these analyses.

### Data and Resource Availability

All raw USV data, processed statistical outputs, and custom MATLAB and Excel-based scripts for post-processing are available from the corresponding author upon reasonable request. USVs were analyzed using Metris Sonotrack (v1.4.7, Metris B.V., Netherlands).

## ACKNOWLEDGEMENTS

We thank the Fralin Biomedical Research Institute at Virginia Tech Carilion School of Medicine Vivarium staff for their assistance with animal care and the Red Gates Foundation and Virginia Tech for start-up funds awarded to M.E.v.d.H. A.L.F. was supported by the Virginia Tech Carilion School of Medicine Summer Research Fellowship.

## AUTHOR CONTRIBUTIONS STATEMENT

Conceptualization: A.L.F, M.E.v.d.H.; Methodology: M.E.v.d.H., A.L.F., B.L.D., A.M.L.; Software: M.E.v.d.H., A.L.F.; Validation: M.E.v.d.H., A.L.F., J.A.C., A.M.L., B.L.D.; Formal analysis: M.E.v.d.H., A.L.F., J.A.C.; Investigation: A.L.F., J.A.C., A.M.L., B.L.D.; Resources: M.E.v.d.H., A.L.F.; Data Curation: M.E.v.d.H., A.L.F., J.A.C.; Writing—original draft: A.L.F., M.E.v.d.H.; Writing—review and editing: M.E.v.d.H., A.L.F., J.A.C., A.M.L., B.L.D.; Visualization: M.E.v.d.H., A.L.F., J.A.C.; Supervision: M.E.v.d.H.; Project administration: M.E.v.d.H., A.L.F., J.A.C.; Funding acquisition: M.E.v.d.H., A.L.F.

